# Network toxicology, AlphaFold 3 and molecular docking for analysing the toxicity of heavy metal ions and their organic complexes in environmental pollutants: A case of Cobalt(II) ion & Bis-(2,4-pentanedionato)cobalt(II)

**DOI:** 10.1101/2025.03.07.642058

**Authors:** Zi-Yong Chu, Xue-Jiao Zi

## Abstract

The aim of this study is to advance the application of network toxicology, AlphaFold 3 and molecular docking techniques in the toxicity analysis of heavy metal ions (HMIs) and heavy metal ions organic complexes (HMIOCs) in environmental pollutants. In this study, the potential toxicity and molecular mechanisms of HMIs and HMIOCs in environmental pollutants were effectively investigated using Cobalt(II) ion & Bis-(2,4-pentanedionato)cobalt(II) (Co(II) & BPCo(II)) as examples. It was found that Co(II) & BPCo(II) regulates multiple signaling pathways by modulating hub targets such as APP, NR3C1, ESR1, CASP3, MMP9, and PTGS2. These pathways include Serotonergic synapse, Cocaine addiction, Pathways of neurodegeneration-multiple diseases, Calcium signaling pathway, Neuroactive ligand-receptor interactions, Alzheimer disease and other pathways, which in turn affects the Blood-Brain Barrier (BBB) in order to destroy neuronal cells, ultimately leading to a variety of mental illnesses (MIs) and nervous system diseases (NSDs), such as Schizophrenia, Depressive disorder, nervous system disorder and Alzheimer Disease, Late Onset. In this study, the toxic effects and molecular mechanisms of Co(II) and BPCo(II) were elucidated, while the limitations of conventional toxicological methods—such as safety issues, time constraints, high expenses, ethical concerns regarding animal use, and limited predictive power—were effectively addressed. These results contribute to the foundation for studying disease diagnosis associated with environmental exposure to HMIs and HMIOCs.

**Graphical Abstract:** 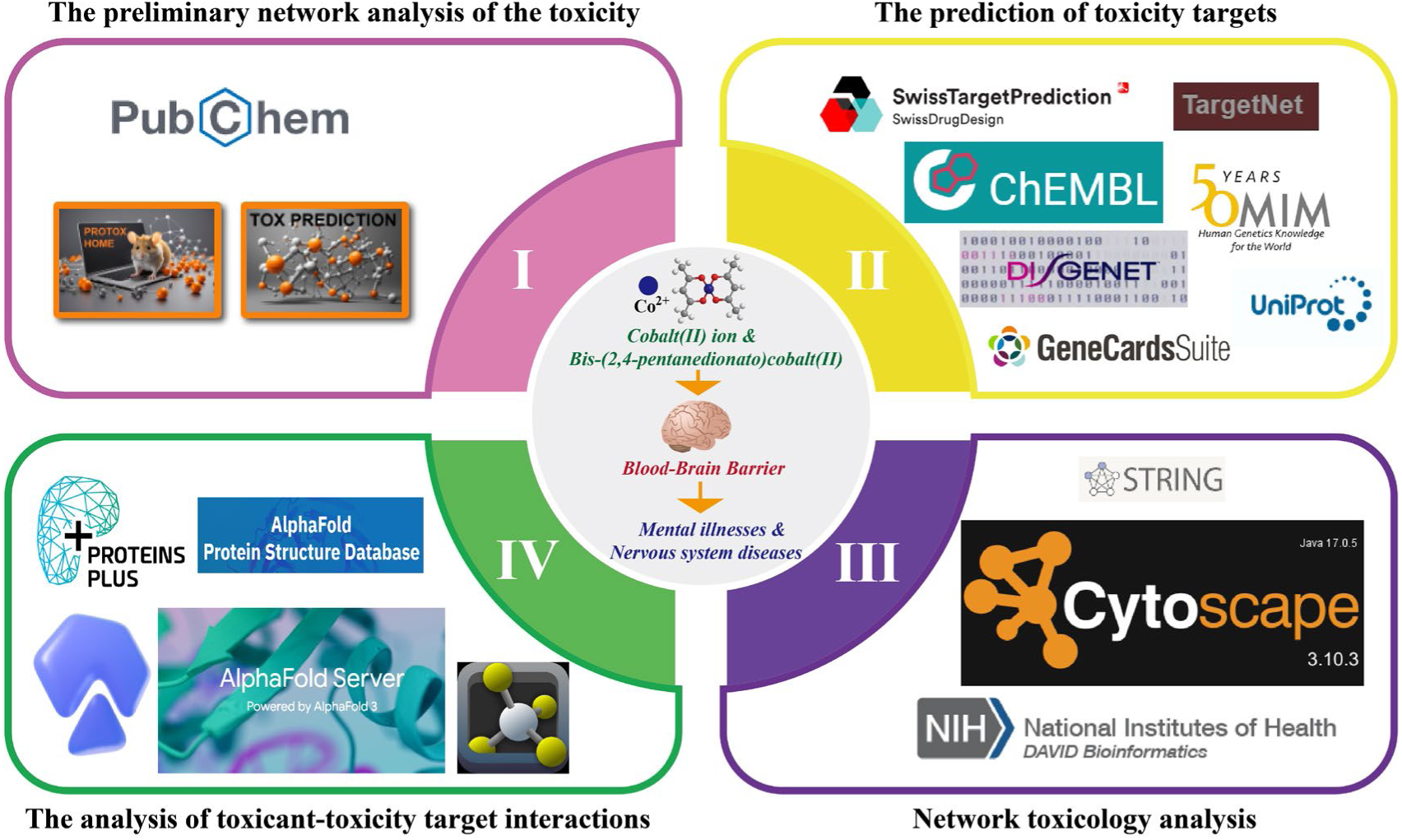

## 1. Introduction

Environmental pollutants are substances that enter the environment as a result of human activities and alter the normal composition and properties of the environment in a way that is directly or indirectly harmful to organisms and humans^1^. Environmental pollutants are classified as chemical, biological, and physical pollutants^2^. With the proliferation of production and use of industrial chemicals, a large number of chemical pollutants with largely unknown toxicological profiles have emerged in the environment^3^. Chemical pollutants are mainly those caused by agrochemicals, food additives, food packaging containers and industrial waste, mercury, cadmium, lead, cyanide, organic phosphorus and other organic or inorganic compounds^4^. Among them, heavy metals (HMs) are hazardous environmental pollutants, which have received widespread attention because they are extremely harmful to human health. The HMs, including lead, cadmium, mercury, arsenic, cobalt and chromium, etc., are found in a wide variety of environments, such as water, air and soil^5^. In addition, the majority of organic pollutants contain electron-rich groups that can act as electron donors to complex with HMIs when coexisting with them^6^. Therefore, assessing the toxicity and molecular mechanisms of HMIs and HMIOCs in environmental pollutants is essential for the prevention and treatment of diseases associated with HMIs and HMIOCs.

HMIs and HMIOCs in environmental pollutants are characterized by a wide range of sources and hazards as well as difficulties in treatment^7^. There are HMIs and HMIOCs that are even radioactive, posing a great threat to human health. Traditional toxicological analyses require the use of animal testing, which is not only time consuming and costly, but may also pose a threat to the health of the researchers themselves^3^. Therefore, there is a need for a safe, efficient, accurate and comprehensive method for the analysis of toxicity and molecular mechanisms of HMIs and HMIOCs in environmental pollutants. Network pharmacology was initially used to study network interactions between drugs and disease targets^8^. The network toxicology was derived on the basis of network pharmacology. Currently, network toxicology has become a hot spot in toxicology research, and its main application is the toxicity and molecular mechanism research of drugs, organic small molecule environmental pollutants and organic small molecule contaminants in food^9–11^. For instance, Huang investigated the toxicity and molecular mechanism of the environmental contaminant Acetyl Tributyl Citrate by network toxicology^3^. Chu et al. investigated the toxicity and molecular mechanism of the food contaminant Aflatoxin B_1_ by network toxicology^9^. However, these studies have been limited to the toxicity and molecular mechanisms of small molecule organic pollutants, and few studies have been reported on the toxicity and molecular mechanisms of HMIs and HMIOCs using network toxicology.

In toxicological studies of chemical pollutants in the environment, the analysis of interactions between chemical pollutants and toxicity targets is important for understanding the toxicity and molecular mechanisms of chemical pollutants. However, current common molecular docking tools such as Autodock Vina and LeDOCK are difficult to meet the needs of HMIs and toxicity target interaction analysis^3,12^. With the development of big data models, many powerful bioinformatics analysis tools have emerged. One example is AlphaFold 3^13^. AlphaFold 3 can be used not only to predict the spatial structure of nucleic acids and proteins, but also to analyze the interactions between ions and proteins and nucleic acids^13^. At present, AlphaFold 3 predicts the interactions of ions such as Mg^2+^, Zn^2+^, Cl^-^, Ca^2+^, Na^+^, Mn^2+^, K^+^, Fe^3+^, Cu^2+^, and Co^2+^ with proteins and nucleic acids^13^. Therefore, AlphaFold 3 is a powerful tool to study the interaction between HMIs and toxic targets.

The common valence states of cobalt consist of Co(II), Co(Ⅲ), Co(Ⅳ), Co(Ⅴ), and Co(Ⅵ). Co(II) is the divalent cation of the element cobalt, which is more toxic than the other valence states^14^. BPCo(II) is an organic complexes of Co(II), which is extremely toxic, and its indiscriminate release can pollute water resources and pose a health risk to humans^15^. There is growing evidence that long-term exposure to cobalt ions and their complexes can lead to a wide range of diseases, such as respiratory tumors, neurological disorders (hearing and visual impairments), cardiovascular and endocrine defects^14–17^. In this study, we used network toxicology, AlphaFold 3 and molecular docking strategies to gain insights into the mechanisms of toxicity of Co(II) and BPCo(II) affecting the BBB leading to various MIs and NSDs, to elucidate the toxicological profiles of Co(II) and BPCo(II) and to predict their potential toxicity and molecular mechanisms. This investigation establishes an efficient and safe methodological framework for evaluating the biological risks posed by HMIs and HMIOCs. The proposed approach demonstrates significant advantages in overcoming the inherent shortcomings of conventional toxicity evaluation methods, particularly regarding temporal efficiency, economic feasibility, ethical considerations in animal experimentation, and prediction accuracy. Furthermore, this methodological advancement creates a knowledge base essential for developing diagnostic research on pathologies linked to HMIs and HMIOCs exposure.

## 2. Methods

### 2.1. The initial network analysis of the toxicity of Co(II) & BPCo(II)

The structural information of Co(II) & BPCo(II) was obtained from PubChem database (https://pubchem.ncbi.nlm.nih.gov/).Subsequently, the coordination complexes of Co(II) & BPCo(II) were subjected to computational toxicology analysis using the ProTox 3.0 (https://tox.charite.de/protox3/) to conduct preliminary toxicity prediction through the platform’s integrated network-based predictive modeling^18,19^.

### 2.2. Targets for collection of Co(II) & BPCo(II)

The structure of the Co(II) was imported into the Index-Calcnet-TargetNet database (http://targetnet.scbdd.com/calcnet/index/)^20^. In addition, the structure of BPCo(II) was entered into the ChEMBL (https://www.ebi.ac.uk/chembl/), SwissTargetPrediction database (http://swisstargetprediction.ch/) and Index-Calcnet-TargetNet (http://targetnet.scbdd.com/calcnet/index/)^20–22^. Subsequently, the forecasts from the databases were transferred to the UniProt database (https://www.uniprot.org/) where “*Homo sapiens*” was chosen for standardization. Following this, the outcomes were consolidated and redundant putative targets were eliminated^23^. Ultimately, the combined targets were utilized to create a target repository for Co(II) & BPCo(II).

### 2.3. Screening for BBB-related targets

We queried GeneCards®: The Human Gene Database (https://www.genecards.org/), DISGENET: Apply for Academic Access (https://disgenet.com/), and OMIM® (https://www.omim.org/) databases for related targets with the search phrase “Blood-Brain Barrier”^24–26^. Furthermore, Venn diagram was employed to identify shared potential targets among Co(II) & BPCo(II) and the BBB. The overlapping section was regarded as a potential target through which Co(II) & BPCo(II) could influence the BBB.

### 2.4. Protein interaction analysis and hub targets screening

The potential targets of Co(II) & BPCo(II) to affect BBB were entered into the STRING 12.0 database (https://cn.string-db.org/), restricting the species to “*Homo sapiens*”, and constructed networks of interactions between potential targets^27^. Outcomes retrieved from the STRING 12.0 database (https://cn.string-db.org/) were subsequently transferred to Cytoscape 3.10.3 for analysis. This software was used to compute the attributes of each node within the network diagram, ultimately constructing a protein-protein interaction (PPI) network diagram^28^.

Subsequently, we applied specific criteria to identify hub targets (those meeting the following conditions were designated as key targets for Co(II) & BPCo(II) affecting the BBB): ① Closeness Centrality exceeds the median, ② Radiality is greater than the median, and ③ Degree value surpasses the median^9^. The MCODE plugin was finally used to screen hub targets for molecular docking and AlphaFold 3 analysis^29^.

### 2.5. Enrichment analysis of gene functions, pathways and diseases of targets

The DAVID database(https://david.ncifcrf.gov/summary.jsp) was used for Gene Ontology (GO) analysis, Kyoto Encyclopedia of Genes and Genomes (KEGG) pathway and diseases enrichment analysis^30^. We elucidated major biological functions through Gene Ontology (GO) analyses, including the assessment of biological processes (BP), cellular components (CC) and molecular functions (MF)^31^. In addition, We investigated the signaling pathways associated with Co(II) & BPCo(II) affecting BBB by KEGG enrichment analysis^32^. We also performed KEGG and diseases enrichment analysis of hub targets of Co(II) & BPCo(II) affecting BBB using the DAVID database. Finally, we visualised the results of the GO, KEGG and diseases enrichment analyses^9^.

### 2.6. The interaction analysis of Co(II) & BPCo(II) and hub targets

The hub targets downloaded from the AlphaFold Protein Structure Database (https://alphafold.ebi.ac.uk/)^33^. Next, we used the ProteinsPlus platform(https://proteins.plus/) to predict the size and location of the molecular docking activity pocket of the hub targets^34^. The AlphaFold 3 (https://golgi.sandbox.google.com/) was utilized to analyze the interactions between Co(II) and hub targets^13^. The AutoDock was employed to analyze the interactions between BPCo(II) and hub targets with reference to the method of Chu et al^12,35^. The analysis was visualized for PyMOL 3.1.

### 2.7. Statistical analysis

The research data was sourced from an online database and was visualized and examined through the SRplot online plotting tool(https://www.bioinformatics.com.cn/)^36^.

## 3. Results

### 3.1. Toxicity analysis of Co(II) & BPCo(II) in an initial network

Using ProTox 3.0 to analyze the toxicity of Co(II) & BPCo(II), we obtained a basic summary of Co(II) & BPCo(II) toxicity(**Fig. 1**). The molecular weight of Co(II) is 58.93 g/mol, and the LD_50_ is 150 mg/Kg. Neurotoxicity toxicity is activated in Organ toxicity, and its Probability is 0.67. Toxicity end points are focused on the BBB-barrier, Ecotoxicity and Nutritional toxicity with Probability of 0.99, 0.73, 0.79 respectively. In addition, the molecular weight of BPCo(II) is 257.15 g/mol, and the LD_50_ is 3200 mg/Kg. Organ toxicity is not activated in BPCo(II), and the toxicity end points are mainly focused on the BBB-barrier, and its Probability is 0.86. In summary, the toxicity end points of Co(II) & BPCo(II) are mainly focused on the BBB-barrier, and these findings provide a basis for further systematic and in-depth studies of the BBB-barrier toxicity effects of Co(II) & BPCo(II) in humans.

**Fig. 1.**
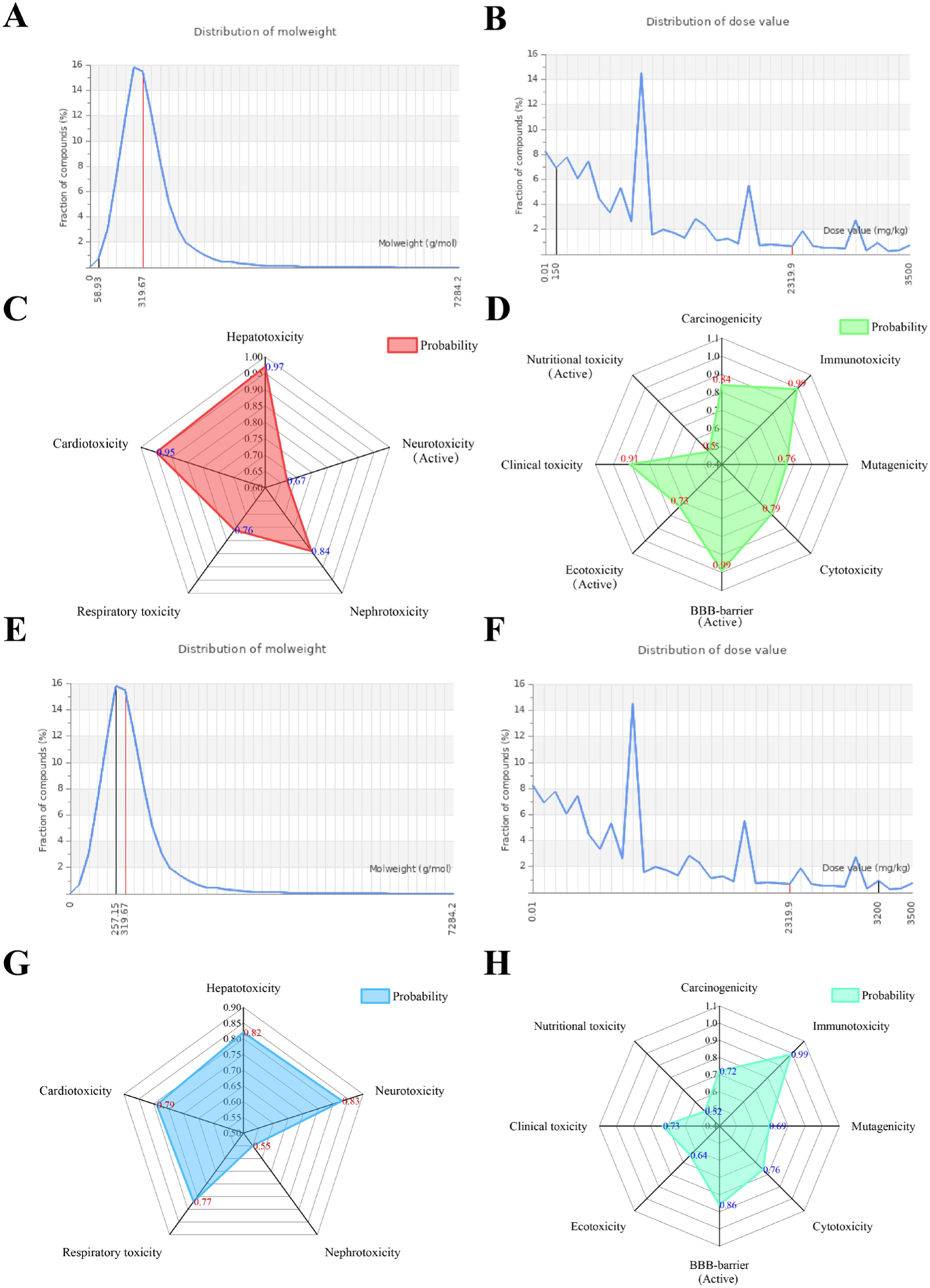
Preliminary toxicity analysis of Co(II) & BPCo(II). (A) The molecular weight of Co(II). (B) The LD50 of Co(II). (C) Radar plot representing the toxicological impact on organs by Co(II). (D) Radar plot of the toxicity end points of Co(II). (E) The molecular weight of BPCo(II). (F) The LD50 of BPCo(II). (G) Radar plot representing the toxicological impact on organs by BPCo(II). (H) Radar plot of the toxicity end points of BPCo(II).

### 3.2. Target of Co(II) & BPCo(II) to affect BBB identified

In this study, we conducted a preliminary screening of 657 Co(II)-related targets using the Index-Calcnet-TargetNet database. Additionally, 523 BPCo(II) targets were identified from the ChEMBL, SwissTargetPrediction, and Index-Calcnet-TargetNet databases. Through GeneCards®: The Human Gene Database, DISGENET: Apply for Academic Access, and OMIM®, we further pinpointed 2365 targets strongly linked to BBB-barrier toxicity. After integrating and removing duplicates from these target sets, we identified 92 overlapping targets, which represent potential candidates for Co(II) & BPCo(II) to influence the BBB(**Fig. 2**).

**Fig. 2.**
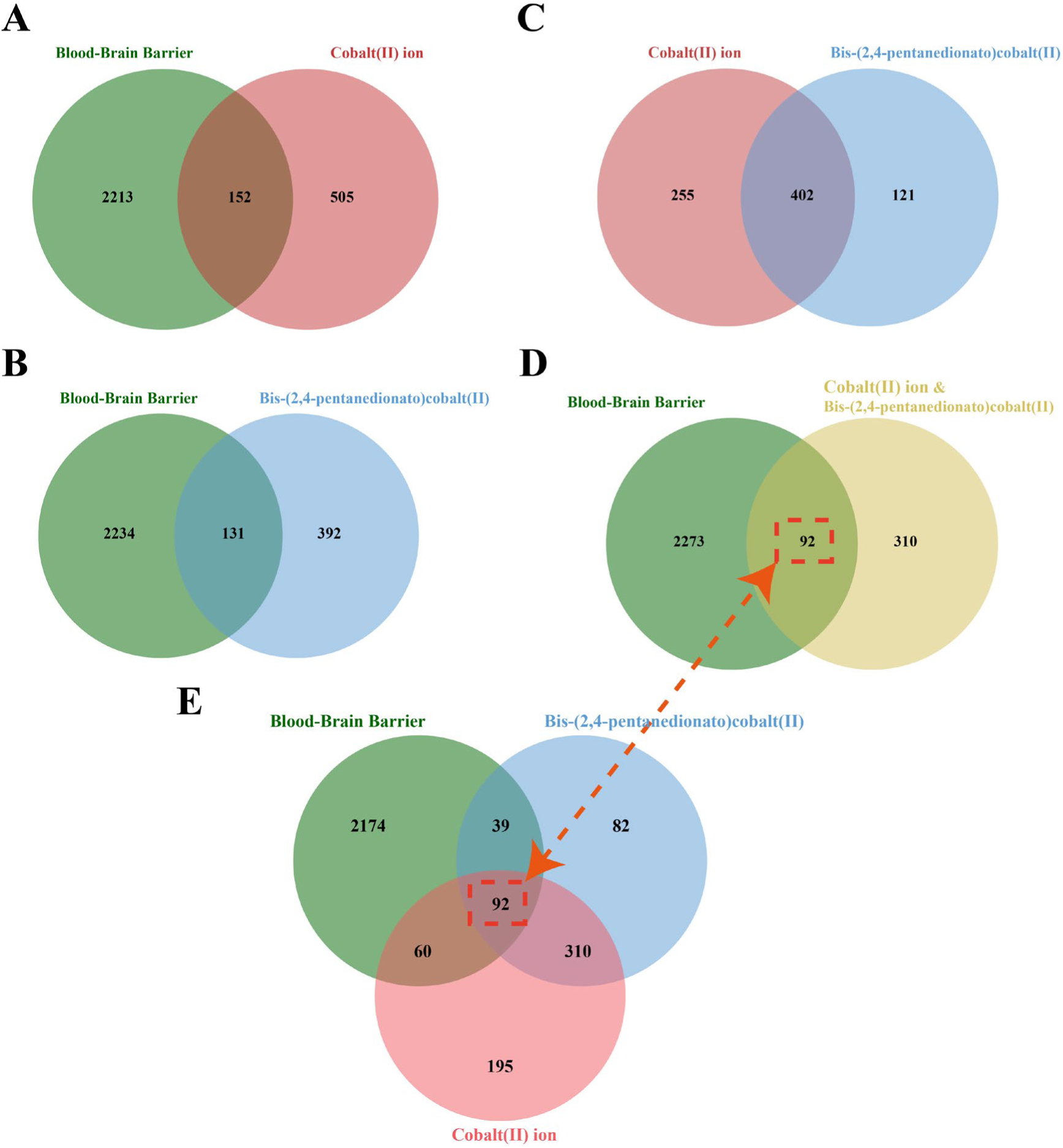
Venn diagram of Co(II) & BPCo(II) and BBB targets

### 3.3. The potential target interaction network and the acquisition of hub targets

Initially, a PPI network comprising 91 nodes and 520 edges was established using the STRING 12.0 database, with an average node degree of 11.4. The network’s topological characteristics, such as Closeness Centrality, Degree, and Radiality, were evaluated using Cytoscape 3.10.3. Additionally, a visual representation of the protein-protein interaction network was created(**Fig. 3**). Subsequently, through network analysis, we pinpointed 39 hub targets of Co(II) & BPCo(II) to affect BBB **(Table S1)**. By employing Cytoscape 3.10.3, we developed a PPI network for these hub targets(**Fig. 4A**), highlighting their specific interactions. We utilized the MCODE plugin to screen six hub targets (Amyloid-beta precursor protein (APP), Glucocorticoid receptor (NR3C1), Estrogen receptor (ESR1), Caspase-3(CASP3), Matrix metalloproteinase-9 (MMP9) and Prostaglandin G/H synthase 2 (PTGS2)) for interaction analysis(**Fig. 4B, Table S2)**. Current research consensus acknowledges that the proteins translated by these genes are crucial in diverse cellular processes, such as governing cell proliferation, survival, and growth; facilitating cellular immune responses; degrading the extracellular matrix (ECM); and mediating cell adhesion and signal transduction. By examining the interplay within this network, the analysis provides a basis for understanding the potential molecular mechanisms by which Co(II) & BPCo(II) may influence the BBB.

**Fig. 3.**
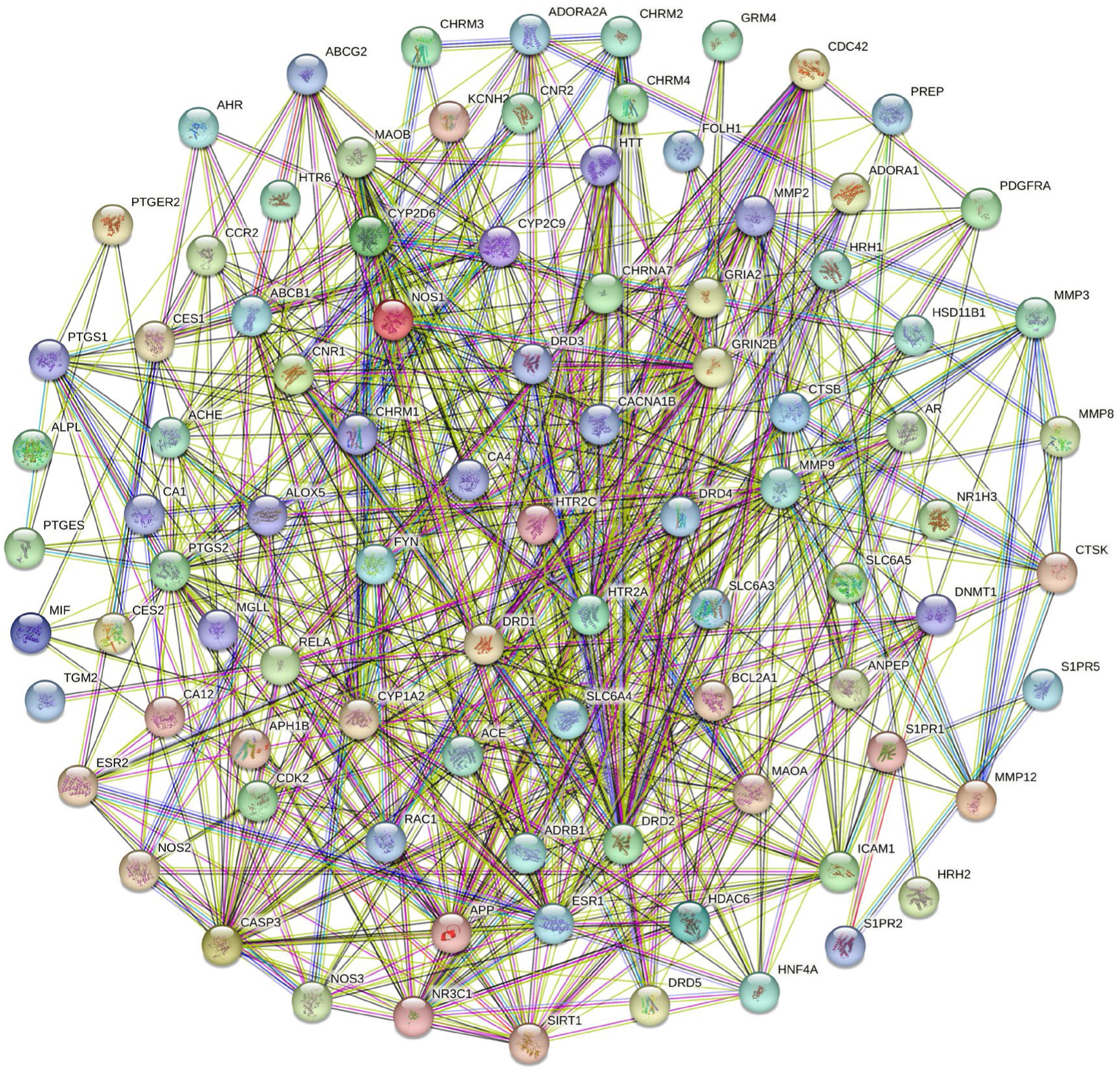
Interaction network diagram of Co(II) & BPCo(II) and BBB potential targets

**Fig. 4.**
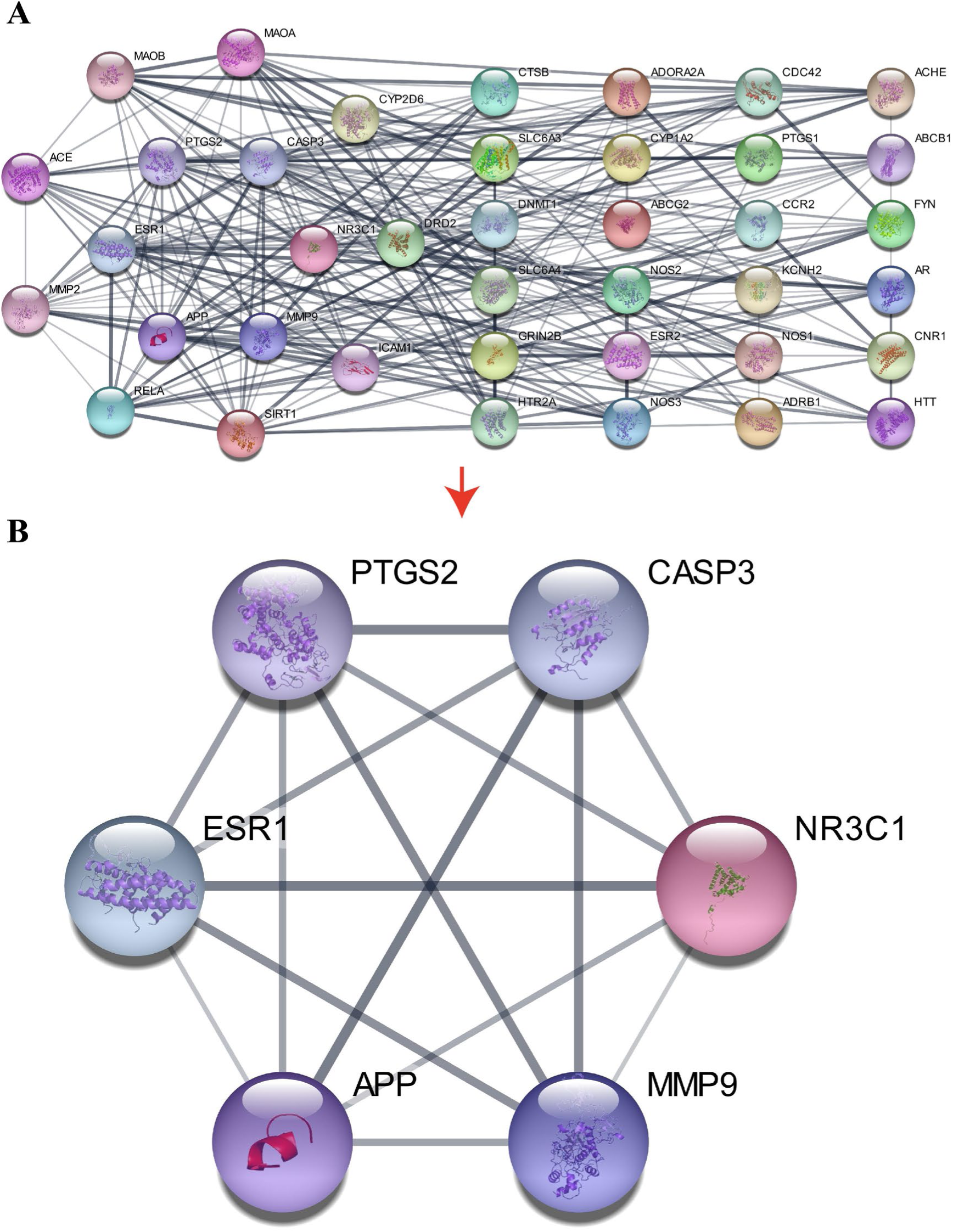
Interaction network diagram of Co(II) & BPCo(II) and BBB hub targets. (A) The hub targets. (B) The hub targets used for molecular docking and AlphaFold 3 analysis.

### 3.4. Functional and pathway enrichment analysis of potential targets

We performed a GO analysis on the 92 potential targets with the DAVID database, focusing on Homo sapiens. This led to the identification of 461 statistically significant GO terms, which included 314 biological processes (BP), 58 cellular components (CC), and 89 molecular functions (MF). The GO terms were arranged based on their false discovery rate (FDR) values, and the top 10 in each category—BP, CC, and MF—with the smallest FDR values are presented(**Fig. 5**). The DAVID database was utilized to conduct KEGG pathway analysis on the 92 potential targets, identifying their involvement in specific signaling pathways. The top 20 KEGG pathways with the smallest FDR values were highlighted among the enriched 60 pathways, visualized through statistical bubble plots and category histograms for significance(**Fig. 6**).

**Fig. 5.**
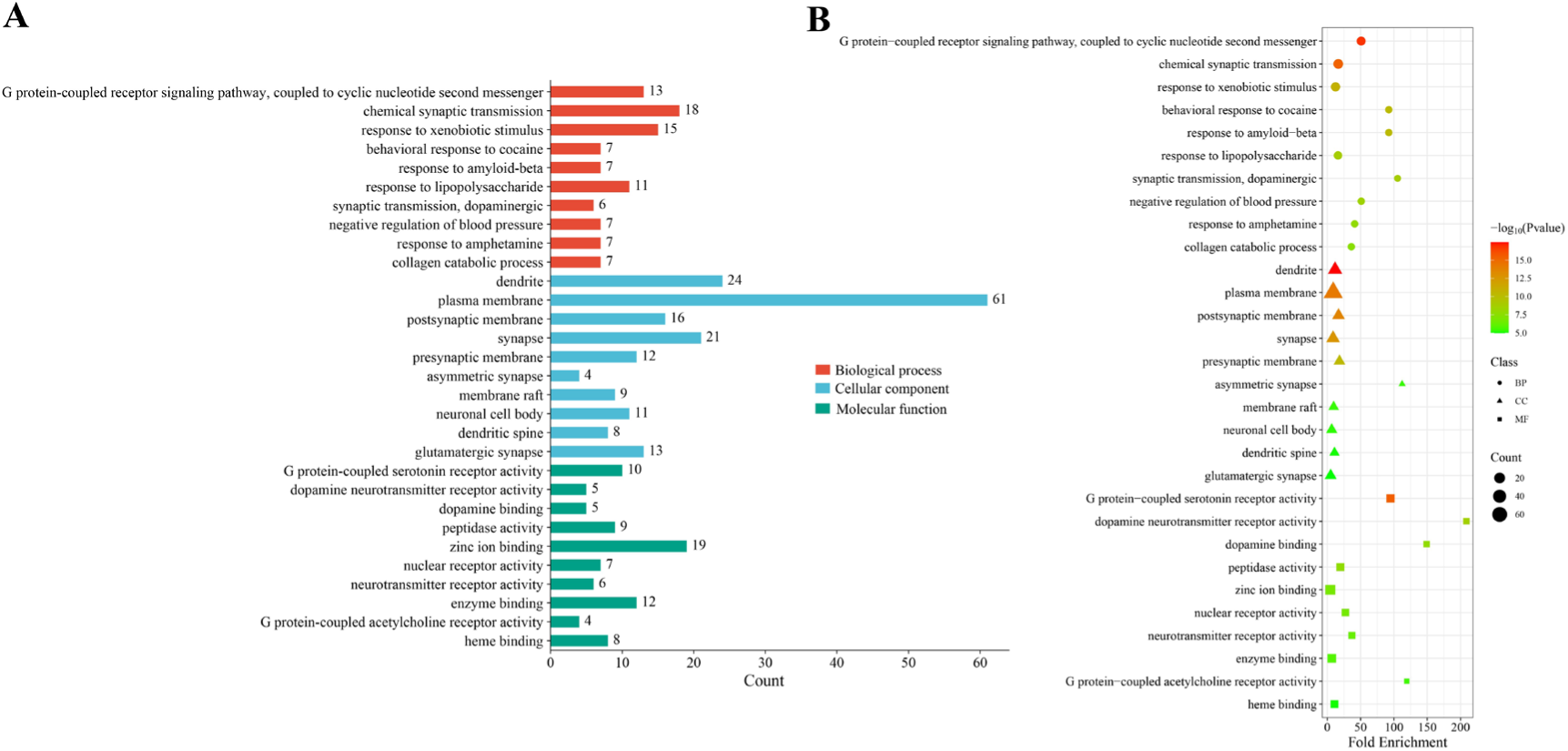
GO Enrichment Analysis for potential targets (Top 10). (A) The bar chart displays the top 10 enrichment results within the GO categories of Biological Process (BP), Cellular Component (CC), and Molecular Function (MF), based on the 92 potential targets with the lowest FDR values. (B) Bubble size is proportional to the level of gene expression within individual pathways.

**Fig. 6.**
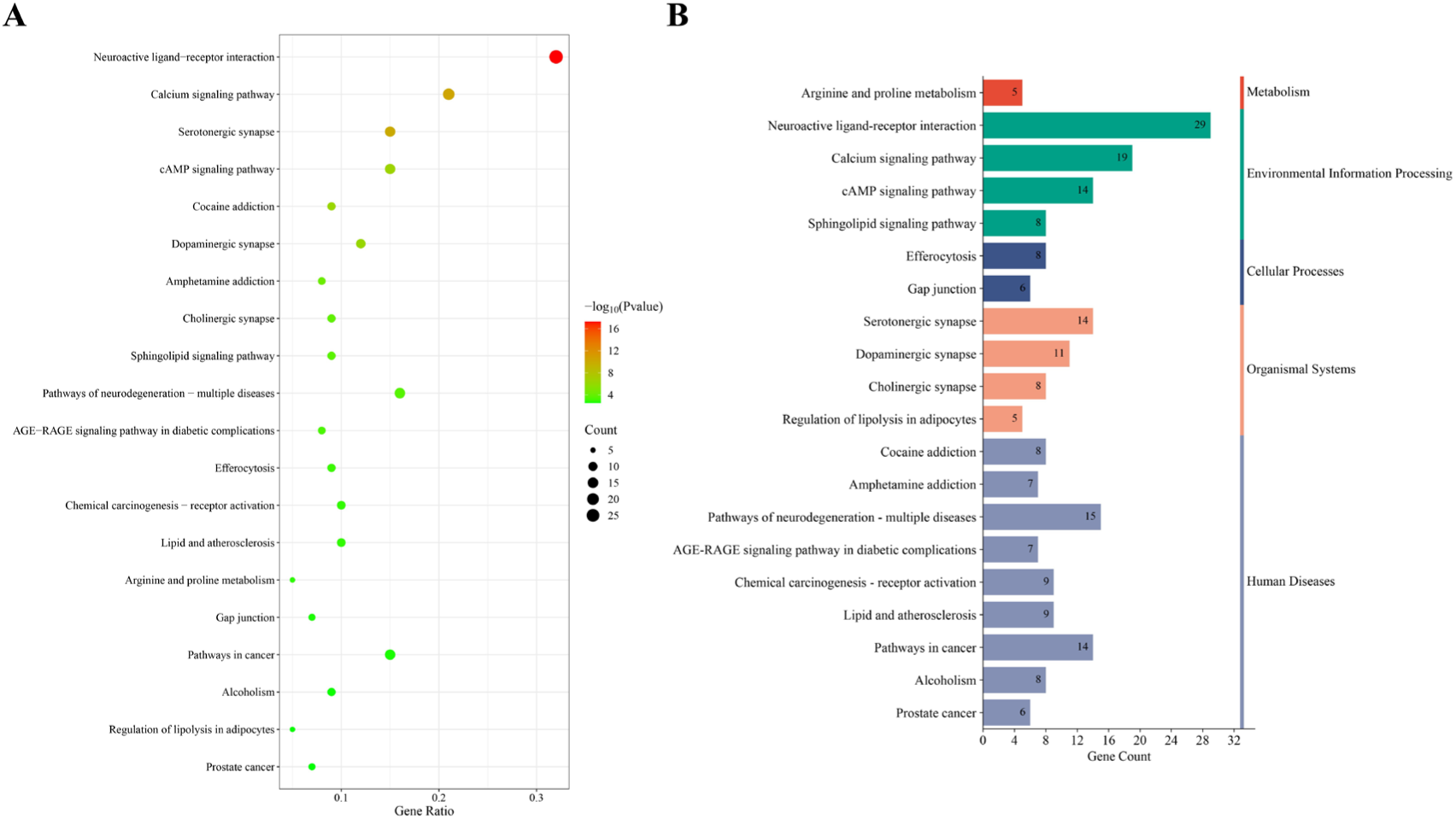
KEGG pathway enrichment analysis for potential targets (Top 20). (A) The bubble chart depicts the top 20 KEGG pathways with enrichment, ranked in descending order by FDR values. (B) The bar graph represents the occurrence and enrichment significance of each pathway.

Significantly, the GO and KEGG analyses of the potential targets revealed that these genes were widely distributed and expressed across various subcellular locations. Additionally, a substantial number of them participated in critical regulatory mechanisms, including the G protein-coupled receptor signaling pathway, coupled to cyclic nucleotide second messenger, chemical synaptic transmission and response to xenobiotic stimulus, etc. It also functions in G protein-coupled serotonin receptor activity, dopamine neurotransmitter receptor activity, zinc ion binding. The enriched KEGG signaling pathways prominently featured those linked to diverse biological processes, such as Neuroactive ligand-receptor interaction, Calcium signaling pathway, Serotonergic synapse, cAMP signaling pathway, Cocaine addiction and other related signalling pathways.

### 3.5. Analysis of pathway and diseases enrichment for hub targets

KEGG and disease enrichment analyses were conducted on 39 identified hub targets of Co(II) & BPCo(II) related to BBB modulation using the DAVID database. A total of 53 signaling pathways were enriched, with the top 25 pathways exhibiting the lowest FDR values selected for visualization and analysis. These were represented through a Sankey diagram illustrating hub target enrichment and a bubble diagram depicting KEGG pathway enrichment for each pathway(**Fig. 7A**). We enriched a total of 791 diseases, selecting the top 30 with the lowest FDR values for visualization and analysis. This included creating a Sankey diagram for hub target enrichment and a bubble diagram for disease enrichment for each of the selected diseases(**Fig. 7B**).

**Fig. 7.**
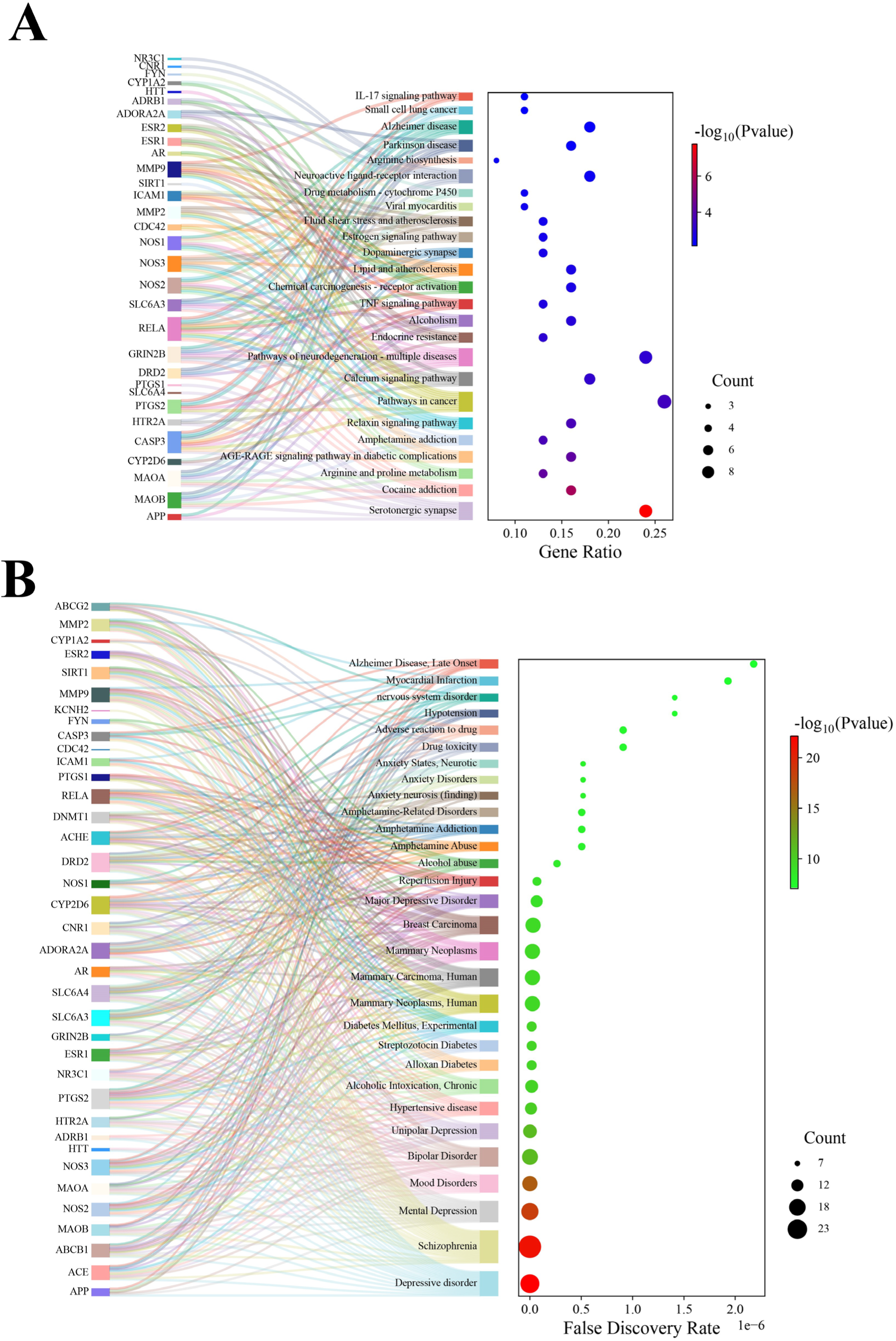
Analysis of pathway and diseases enrichment for hub targets. (A) KEGG pathway enrichment analysis for hub targets Sankey and bubble diagrams (Top 25). (B) Enrichment of diseases for hub targets Sankey and bubble diagrams (Top 30).

The primary pathways involving hub targets linked to Co(II) & BPCo(II) and their impact on the BBB are intricately connected to pathways associated with numerous MIs and NSDs, such as Schizophrenia, Depressive disorder, nervous system disorders, and Alzheimer’s Disease, Late Onset, among others, as indicated in our research findings(**Fig. 7B**). The process encompasses various molecularly-driven signaling pathways, including Serotonergic synapse, Cocaine addiction, Neurodegeneration pathways associated with multiple diseases, Calcium signaling pathway, Neuroactive ligand-receptor interactions, and Alzheimer’s disease(**Fig. 6 and 7A**). Interestingly, there is a link between the influence of Co(II) & BPCo(II) on the BBB and a broad spectrum of diseases, with this association evident for both potential and hub targets(**Figs. 6, 7)**. This findings seem to suggest that Co(II) & BPCo(II) may affect the BBB, leading to a variety of MIs and NSDs, although further research is needed.

### 3.6. The results of AlphaFold 3 analysis of Co(II) with hub targets

To explore the interaction of Co(II) with the six hub targets, we analyzed the interaction of Co(II) with APP, ESR1, NR3C1, CASP3, MMP9 and PTGS2 using AlphaFold 3(**Fig. 8, Fig. S1, Table S3)**. The results showed that Co(II) interacts with negatively charged acidic amino acids, as well as cation-п interactions with the imidazole ring of histidine(His) and the benzene ring of tryptophan (Trp); Co(II) is surrounded by a large number of basic amino acids, namely arginine (Arg) and histidine (His). In addition, S-containing cysteines (Cys) were also present around it. These interactions may be one of the reasons why Co(II) affects the BBB leading to various MIs and NSDs.

**Fig. 8.**
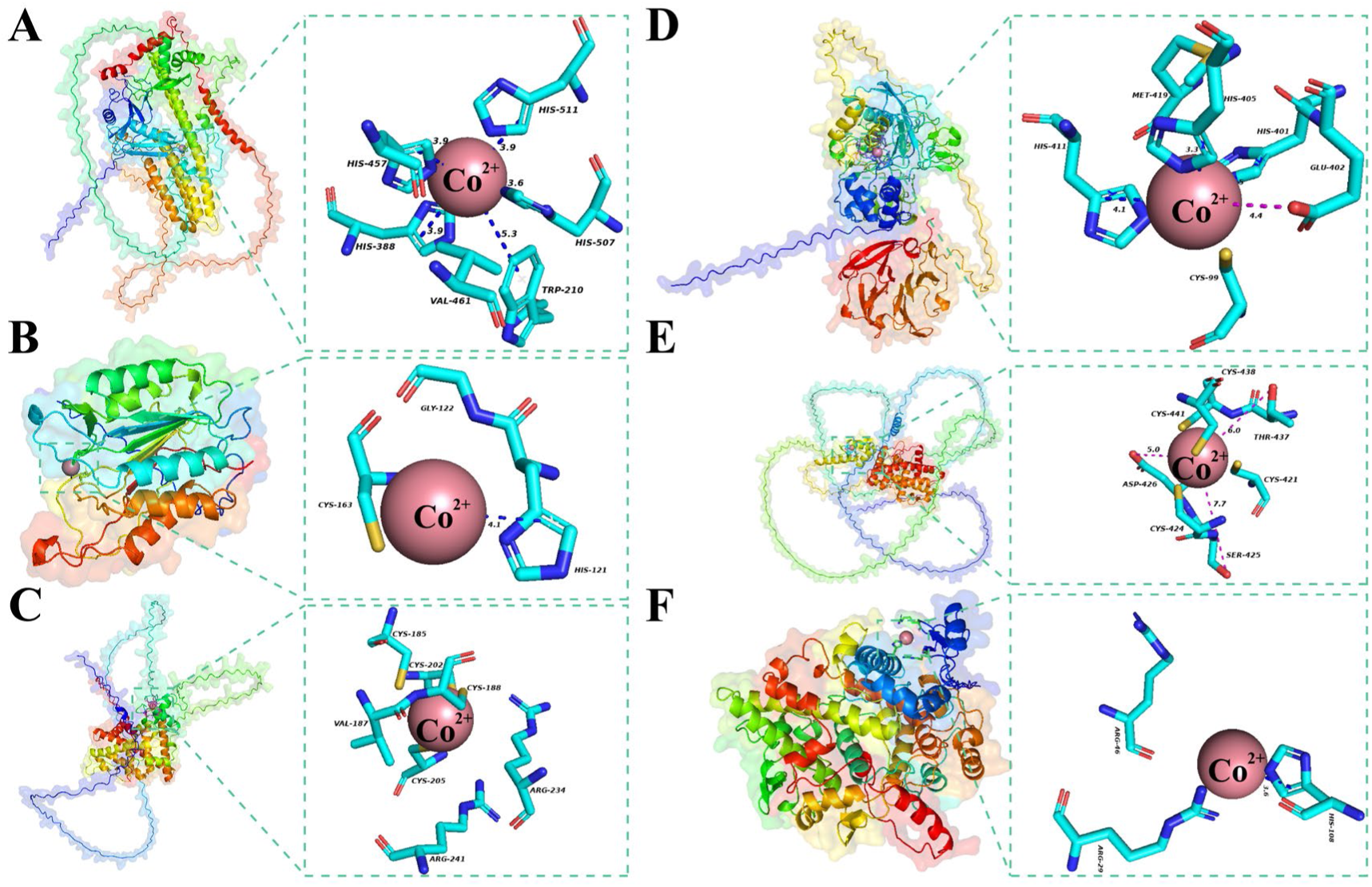
Illustration of AlphaFold 3 analysis outcomes for hub targets. (A) Co(II) and APP, (B) Co(II) and CASP3, (C) Co(II) and ESR1, (D) Co(II) and MMP9, (E) Co(II) and NR3C1, (F) Co(II) and PTGS2.

### 3.7. The results of molecular docking analysis of BPCo(II) with hub targets

A molecular docking study was conducted to examine the interactions between BPCo(II) and six key targets (APP, ESR1, NR3C1, CASP3, MMP9, and PTGS2). The Autodock software was employed to produce six molecular docking outcomes with reduced binding energies. These outcomes revealed a significant attraction between BPCo(II) and the hub targets. Significantly, the binding energies for all six hub targets with BPCo(II) were below −5.5 Kcal/mol, suggesting that BPCo(II) binds spontaneously to each of these targets, indicating their crucial function in the molecular pathway by which BPCo(II) impacts the BBB(**Fig. 9, Fig. S2, Table S4)**. These interactions may be one of the reasons why BPCo(II) affects the BBB leading to various MIs and NSDs.

**Fig. 9.**
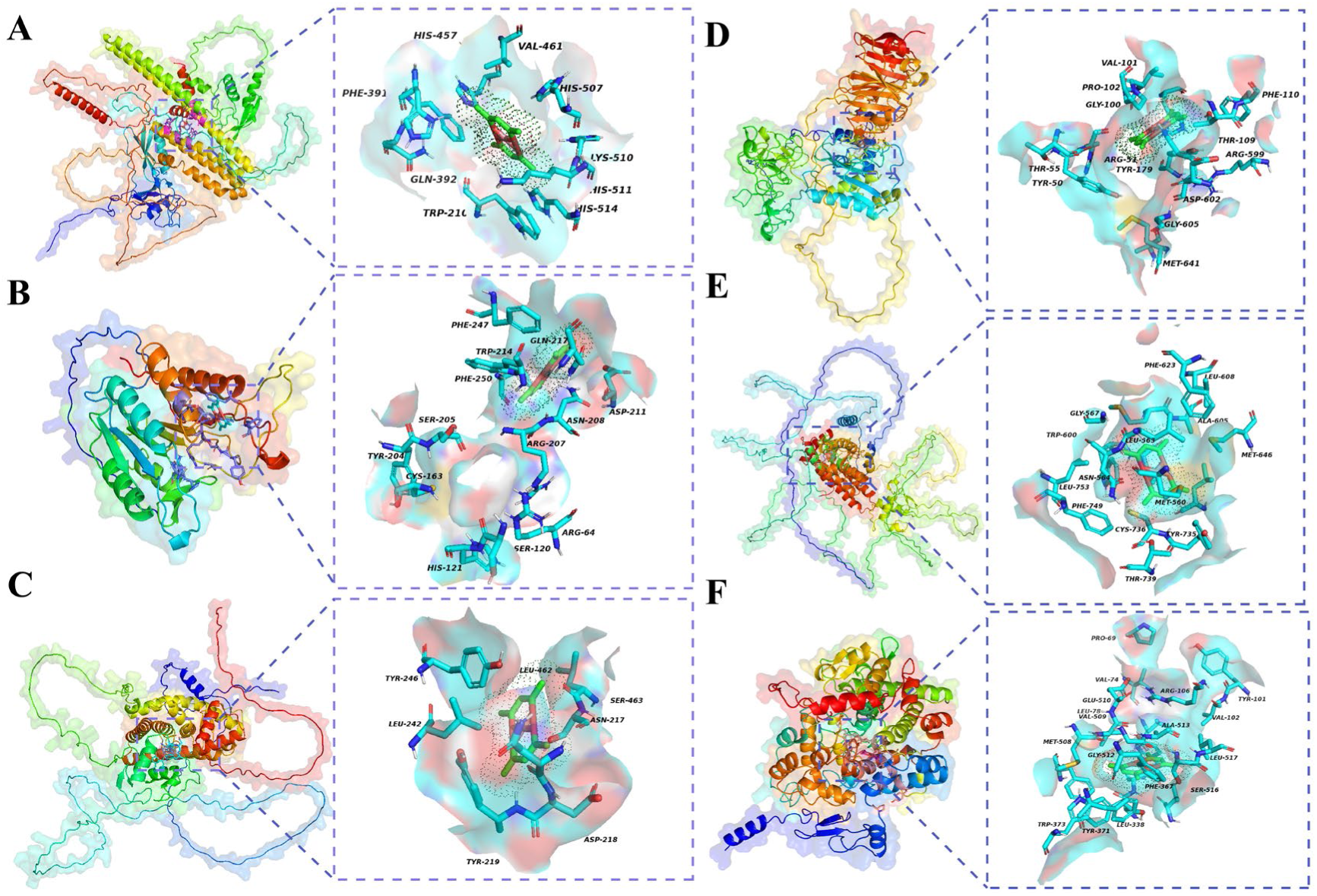
Visualisation of molecular docking results for hub targets. (A) BPCo(II) and APP, (B) BPCo(II) and CASP3, (C) BPCo(II) and ESR1, (D) BPCo(II) and MMP9, (E) BPCo(II) and NR3C1, (F) BPCo(II) and PTGS2.

## 4. Discussion

It is important to analyze the toxicity and molecular mechanism of HMIs and HMIOCs in environmental pollutants, which are very harmful to human beings, for the prevention and treatment of diseases related to HMIs and HMIOCs^5,7^. For this research, Co(II) and BPCo(II) served as case studies. An initial toxicity evaluation of Co(II) and BPCo(II) was carried out for the first time using ProTox 3.0, and it was found that Co(II) & BPCo(II) may have BBB toxicity. The toxicity-related targets, diseases and signaling pathways of Co(II) & BPCo(II) were predicted for the first time using a network toxicology approach. Furthermore, we examined the interactions between Co(II) & BPCo(II) and the respective toxic targets through AlphaFold 3 and molecular docking techniques, aiming to offer novel insights into the toxicity research of HMIs and HMIOCs within environmental pollutants.

The initial toxicity analysis of Co(II) & BPCo(II) showed that Co(II) & BPCo(II) is primarily BBB toxic. This means that Co(II) & BPCo(II) may cause severe damage to the BBB in humans and may destroy nerve cells, thereby increasing the risk of MIs and NSDs. The KEGG enrichment analysis indicated that Co(II) & BPCo(II) may be involved in the Neuroactive ligand-receptor interaction pathway. It is proposed that Co(II) & BPCo(II) may negatively affect the nervous system by affecting the BBB, leading to various MIs and NSDs, such as Schizophrenia, Depressive disorder, nervous system disorder and Alzheimer Disease, Late Onset. Hence, it remains essential to enhance the toxicity evaluation of Co(II) & BPCo(II) for a more thorough comprehension of their potential risks to human health.

This research identified 39 key targets linked to Co(II) & BPCo(II) that contribute to multiple MIs and NSDs by impacting the BBB, including APP, NR3C1, ESR1, CASP3, MMP9, and PTGS2. APP is a transmembrane protein that is highly expressed in brain tissues and plays an important role in cell signaling, synapse formation and other physiological processes^37^. In vitro experiments have shown that the complex formed by the combination of Cu^2+^ and APP may cause changes in the structure or function of APP, which may produce toxic effects on neurons and lead to neuronal death^37,38^. In addition, Cu^2+^ have oxidizing properties, which can promote the oxidation of LDL, and the oxidized LDL may trigger a series of intracellular reactions, which further exacerbate the toxic effects of APP on neurons and worsen neuronal death^37,38^. It is worth noting that Cu^2+^ is a divalent cation and BPCo(II) contains divalent Co, which is itself a divalent cation. Therefore, we hypothesize that Co(II) & BPCo(II) can also cross the BBB and cause neuronal damage by the similar mechanism as Cu^2+^, leading to various MIs and NSDs.

NR3C1 is a receptor protein capable of binding to glucocorticoids. The glucocorticoids are an important class of steroid hormones with a variety of physiological functions, such as participating in the regulation of inflammatory responses, cell proliferation and differentiation, and other processes^39^. It was shown that activation of the NR3C1 may upregulate miR-210-3p and miR-362-5p, two miRNAs that target and inhibit the SHH signaling pathway, leading to impaired axonal development^40^. Therefore, we hypothesize that Co(II) & BPCo(II) may affect the development of the nervous system by crossing the BBB, leading to some MIs and NSDs.

The ESR1 is a class of receptor proteins located in the nucleus of the cell, which can bind to the corresponding hormone ligands and thus regulate the transcription process of genes, which in turn affects a variety of biological functions, such as cell proliferation, differentiation, metabolism, and so on^41,42^. In the presence of circulating estradiol (17-beta-estradiol/E2), ESR1 maintains neuronal survival in ischemia-reperfusion injury^43^. ESR1’s significant involvement in the neuronal damage caused by Co(II) & BPCo(II) traversing the BBB, resulting in various MIs and NSDs, is evident, and this aligns with the observations reported in the literature.

CASP3, a pivotal enzyme in the apoptosis pathway, is implicated in the management of conditions including cancer, heart failure, and neurodegenerative disorders^44^. It was shown that CoCl₂ exposure led to excessive accumulation of ROS in the cytoplasm and mitochondria of SH-SY5Y cells, which resulted in apoptosis^45^. In addition, CoCl₂-induced hypoxia not only activated the endogenous apoptotic pathway, but also induced CASP3/GSDME-dependent and NLRP3/CASP1/GSDMD-mediated focal death^45^. This suggests that two programmed cell death pathways, apoptosis and pyroptosis, were initiated simultaneously in neuronal cells under CoCl₂ exposure, and that the activation of the pyroptosis pathway was closely related to the expression of GSDM^45^. It is noteworthy that cobalt ions in CoCl₂ belong to the same class of Co(II) as Co(II) & BPCo(II). Therefore, we hypothesize that Co(II) & BPCo(II) and CoCl₂ cause focalization or apoptosis of neuronal cells through the BBB by similar molecular mechanisms, which leads to a variety of MIs and NSDs.

The MMP9 is an enzyme capable of degrading the extracellular matrix and plays an important role in a variety of physiological and pathological processes, including tissue remodeling, inflammatory response, tumor invasion and metastasis^46^. It has been demonstrated that exposure to arsenic (As) causes MMP2 and MMP9-mediated BBB disruption as well as neuronal apoptosis, leading to cognitive dysfunction^47^. As and Co(II) & BPCo(II) are widespread environmental pollutants with similar chemical properties ^16,47^. Therefore, we hypothesize that Co(II) & BPCo(II) may lead to apoptosis of neuronal cells by destroying the BBB, thereby producing MIs and NSDs.

The PTGS2, often abbreviated as PGHS-2 or COX-2, is the main isoenzyme responsible for the production of inflammatory prostaglandins^48^. It was shown that Doxorubicin accelerated cognitive decline in rats by increasing neuroinflammation, oxidative stress, apoptosis, and mitochondrial activity through increased levels of NF-κB and COX-2, decreased levels of SOD, and increased levels of Bax, CASP3, and lipid peroxidation^49^. It was also found that inhibition of COX-2 protects zebrafish from PTZ-induced neurological damage and behavioral changes by attenuating the inflammatory response^50^. This shows that PTGS2 has an important role in nerve injury-related diseases. Hence, we hypothesized that Co(II) and BPCo(II) may induce neuronal apoptosis by disrupting the BBB, thereby triggering a number of MIs and NSDs associated with neuronal cell damage.

The results of functional, disease and pathway enrichment analyses of potential and hub targets suggest that Co(II) and BPCo(II) may affect the BBB by modulating pathways such as Serotonergic synapse, Cocaine addiction, Pathways of neurodegeneration-multiple diseases, Calcium signaling pathway, Neuroactive ligand-receptor interactions, Alzheimer disease and other pathways destroying neuronal cells, leading to a variety of MIs and NSDs, such as Schizophrenia, Depressive disorder and Alzheimer Disease, Late Onset. However, further studies are needed to verify this. AlphaFold 3 analysis showed that there were cation-Π, salt bridge and other interactions between the six hub targets and Co(II). The molecular docking outcomes revealed that the six key targets binds stably to BPCo(II) with binding energies below −5.5 Kcal/mol. Thus, it is clear that the six hub targets play an important role in the effect of BPCo(II) on the BBB to rupture neuronal cells, which leads to a variety of MIs and NSDs.

In addition to providing insights into the molecular mechanisms by which Co(II) & BPCo(II) affects the BBB leading to various MIs and NSDs, this study proposes strategies for network toxicology, AlphaFold 3 and molecular docking for the safe and rapid study of toxicity and molecular mechanisms of HMIs and HMIOCs in the environment. We performed the first target prediction of HMIs and HMIOCs using Index-Calcnet-TargetNet. The AlphaFold 3 enables high accuracy prediction of biomolecular structures and interactions^13^. For the first time, we realized the interaction analysis between HMIs and toxic targets using AlphaFold 3. Unlike conventional environmental toxicology methods, our approach involves analyzing the toxicity of HMIs and HMIOCs in environmental pollutants using network toxicology, AlphaFold 3, and molecular docking. This methodology achieves a safer and more efficient environmental toxicology analysis, while also reducing the risk to experimenters during the experimental phase. Concurrently, it facilitates large-scale data handling, incorporates various influencing factors, and enables the exploration of multi-target, multi-pathway, and multi-disease mechanisms, as well as the prediction of novel toxicities. This approach also minimizes and prevents the ethical issues associated with animal testing.

The promising findings notwithstanding, we recognize the need for targeted experimental validation to confirm the results of toxicity analyses of HMIs and HMIOCs. Future studies should include large-scale epidemiological analyses in conjunction with relevant animal models to identify identified central targets, pathways and diseases. This will ultimately elucidate preventive and therapeutic strategies for diseases associated with HMIs and HMIOCs in environmental pollutants. This study will provide a solid theoretical basis for preventive and therapeutic strategies for Co(II) & BPCo(II) affecting the BBB and destroying nerve cells leading to various MIs and NSDs and other health risks associated with HMIs and their complexes.

## 5. Conclusions

In this paper, the targets and mechanisms of the toxic effects of Co(II) & BPCo(II) in environmental pollutants were investigated based on network toxicology, AlphaFold 3 and molecular docking techniques. A total of 39 hub targets of Co(II) & BPCo(II) were identified. Co(II) & BPCo(II) can regulate signaling pathways such as Serotonergic synapse, Cocaine addiction, Pathways of neurodegeneration-multiple diseases, Calcium signaling pathway, Neuroactive ligand-receptor interactions, Alzheimer disease and other pathways through hub targets such as APP, NR3C1, ESR1, CASP3, MMP9, and PTGS2, which affects the BBB leading to a variety of MIs and NSDs, such as Schizophrenia, Depressive disorder, nervous system disorder and Alzheimer Disease, Late Onset. This research not only clarified the potential toxicity and molecular mechanisms of Co(II) & BPCo(II) but also introduced a novel strategy employing network toxicology, AlphaFold 3, and molecular docking to analyze the toxicity and molecular biological mechanisms of HMIs and HMIOCs within environmental pollutants. This approach surmounts the constraints pertaining to safety, efficiency, expense, and ethical concerns associated with animal experimentation, as well as the predictive limitations of conventional toxicological evaluations. Additionally, it establishes a groundwork for diagnostic research on diseases linked to exposure to HMIs and HMIOCs in environmental contaminants.

## CRediT authorship contribution statement

Zi-yong Chu designed the experiment and performed the statistical analysis. Zi-yong Chu and Xue-Jiao Zi carried out the study. Zi-yong Chu and Xue-Jiao Zi drafted the manuscript. All authors read and approved the final manuscript.

## Acknowledgement

The authors declare that they have no known competing financial interests or personal relationships that could have appeared to influence the work reported in this paper. This research did not receive any specific grant from funding agencies in the public, commercial, or not-for-profit sectors.

## Data availability

Data will be made available on request.

---

**Table S1.** hub targets screened from PPI network.

| Gene | Radiality | ClosenessCentrality | Degree | Gene | Radiality | ClosenessCentrality | Degree |
| --- | --- | --- | --- | --- | --- | --- | --- |
| ABCB1 | 0.7627 | 0.9961 | 62 | DNMT1 | 0.6767 | 0.9940 | 47 |
| SIRT1 | 0.7317 | 0.9954 | 57 | PTGS1 | 0.7500 | 0.9958 | 60 |
| MMP2 | 0.7087 | 0.9949 | 53 | DRD2 | 0.7377 | 0.9956 | 58 |
| NR3C1 | 0.8108 | 0.9971 | 69 | PTGS2 | 0.9000 | 0.9986 | 80 |
| SLC6A4 | 0.7258 | 0.9953 | 56 | ADRB1 | 0.6716 | 0.9939 | 46 |
| KCNH2 | 0.6667 | 0.9938 | 45 | MMP9 | 0.8182 | 0.9972 | 70 |
| ICAM1 | 0.7377 | 0.9956 | 58 | AR | 0.7200 | 0.9951 | 55 |
| SLC6A3 | 0.7143 | 0.9950 | 54 | MAOB | 0.7895 | 0.9967 | 66 |
| APP | 0.8738 | 0.9982 | 77 | RELA | 0.6569 | 0.9935 | 43 |
| ACE | 0.7895 | 0.9967 | 66 | ESR1 | 0.8824 | 0.9983 | 78 |
| CCR2 | 0.6716 | 0.9939 | 46 | HTR2A | 0.6667 | 0.9938 | 45 |
| NOS3 | 0.7317 | 0.9954 | 57 | NOS1 | 0.6569 | 0.9935 | 43 |
| ACHE | 0.7895 | 0.9967 | 66 | ADORA2A | 0.7200 | 0.9951 | 55 |
| CASP3 | 0.8411 | 0.9976 | 73 | CDC42 | 0.6618 | 0.9936 | 44 |
| NOS2 | 0.6716 | 0.9939 | 46 | ABCG2 | 0.7031 | 0.9947 | 53 |
| MAOA | 0.7826 | 0.9965 | 65 | CYP2D6 | 0.6870 | 0.9943 | 49 |
| CYP1A2 | 0.6923 | 0.9944 | 50 | HTT | 0.6716 | 0.9939 | 46 |
| ESR2 | 0.7563 | 0.9960 | 61 | CNR1 | 0.6767 | 0.9940 | 47 |
| CTSB | 0.6818 | 0.9942 | 49 | GRIN2B | 0.7031 | 0.9947 | 52 |
| FYN | 0.6923 | 0.9944 | 50 | **Median** | **0.6536** | **0.9931** | **40.6813** |

**Table S2.** Molecular docking parameters for hub targets.

| Gene/ Protein | Protein | Uniprot ID | X | Y | Z |
| --- | --- | --- | --- | --- | --- |
| APP | Amyloid-beta precursor protein | P05067 | -17.11 | 0.57 | -7.94 |
| NR3C1 | Glucocorticoid receptor | P04150 | -1.45 | 4.18 | -3.61 |
| CASP3 | Caspase-3 | P42574 | 14.58 | 6.01 | -6.79 |
| ESR1 | Estrogen receptor | P03372 | 1.69 | -3.80 | 11.81 |
| MMP9 | Matrix metalloproteinase-9 | P14780 | -15.68 | -0.29 | 5.09 |
| PTGS2 | Prostaglandin G/H synthase 2 | P35354 | 6.26 | 8.44 | -2.17 |

**Table S3.** The results of the AlphaFold 3 analysis of hub targets with Co(II)

| Gene/ Protein | Heavy metal ions | ipTM | pTM |
| --- | --- | --- | --- |
| APP | Co(II) | 0.96 | 0.34 |
| NR3C1 | Co(II) | 0.81 | 0.38 |
| CASP3 | Co(II) | 0.73 | 0.82 |
| ESR1 | Co(II) | 0.74 | 0.46 |
| MMP9 | Co(II) | 0.84 | 0.5 |
| PTGS2 | Co(II) | 0.76 | 0.9 |

**Table S4.** Molecular docking results for hub targets and BPCo(II)

| Gene/ Protein | Affinity (Kcal/mol) | Min RMSD | Binding of amino acid residues |
| --- | --- | --- | --- |
| APP | -6.1 | 0 | His457, Val461, His570, Lys510, His511, His514, Trp210, Glu392, Phe391 |
| NR3C1 | -5.9 | 0 | Phe623, Leu608, Ala605, Met646, Met560, Trp735, Cys736, Thr739, Phe749, Leu753, Asn564, Trp600, Gly567 |
| CASP3 | -5.5 | 0 | Gln217, Asp211, Asn208, Arg207, Arg64, Ser1220, His121, Cys163, Trp204, Ser205, Phe250, Trp214, Phe247 |
| ESR1 | -6.7 | 0 | Tyr246, Asn217, Leu242, Trp19, Asp218 |
| MMP9 | -6.3 | 0 | Phe110, Thr109, Arg599, Asp602, Arg51, Tyr50, Thr55, Pro102, Gly100, Val101, Lys603, Gly605, Met641, Arg51, Tyr179 |
| PTGS2 | -6.9 | 0 | Leu78, Tyr101, Val102, Leu517, Ser516, Phe367, Leu338, Tyr371, Trp373, Met508, Val509, Glu510, Val74, Pro69, Val335, Phe504, Gly512, Ala513, Arg106 |

**Fig. S1.**
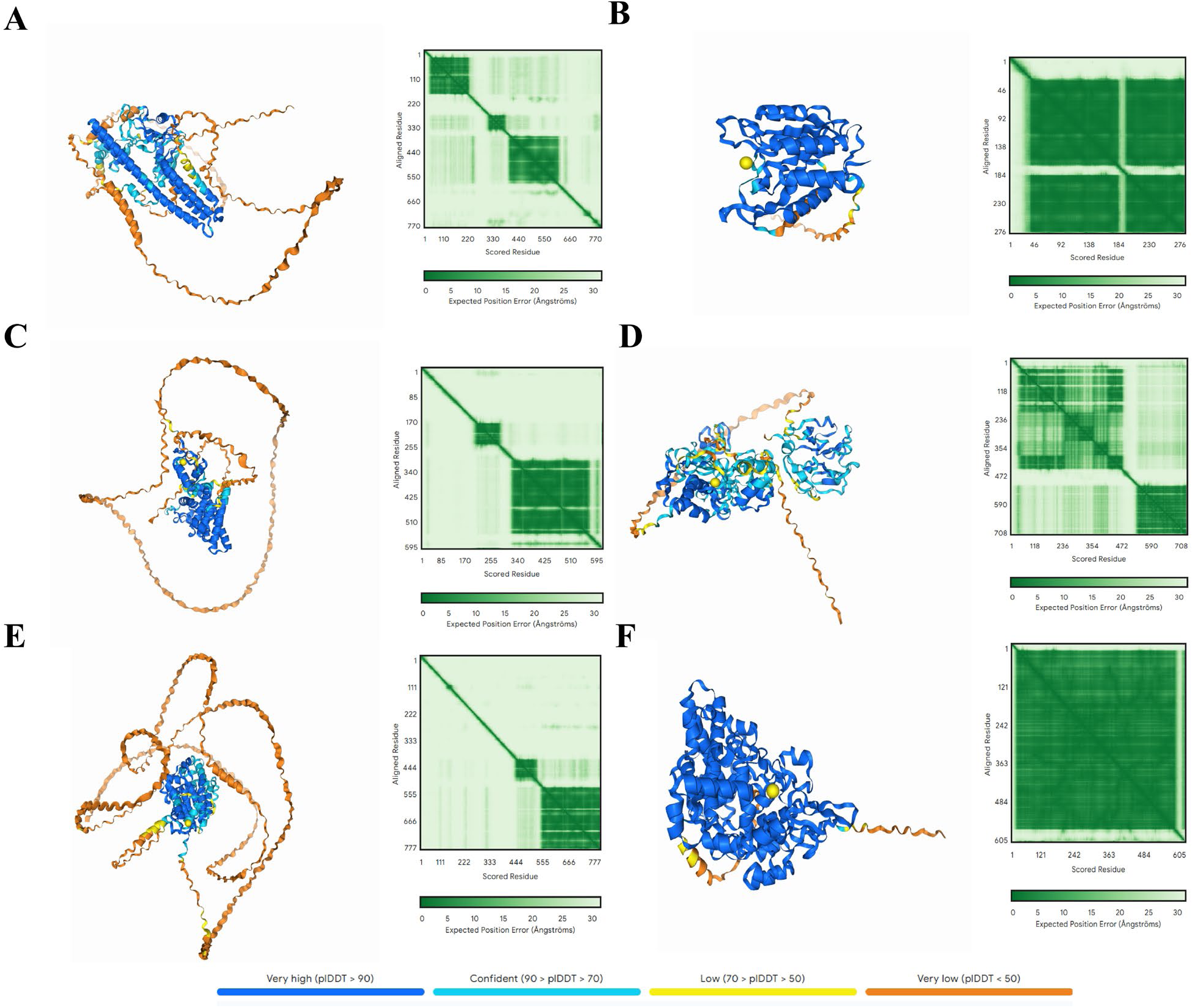
The results of the AlphaFold 3 analysis of hub targets with Co(II). (A) Co(II) and APP, (B) Co(II) and CASP3, (C) Co(II) and ESR1, (D) Co(II) and MMP9, (E) Co(II) and NR3C1, (F) Co(II) and PTGS2.

**Fig. S2.**
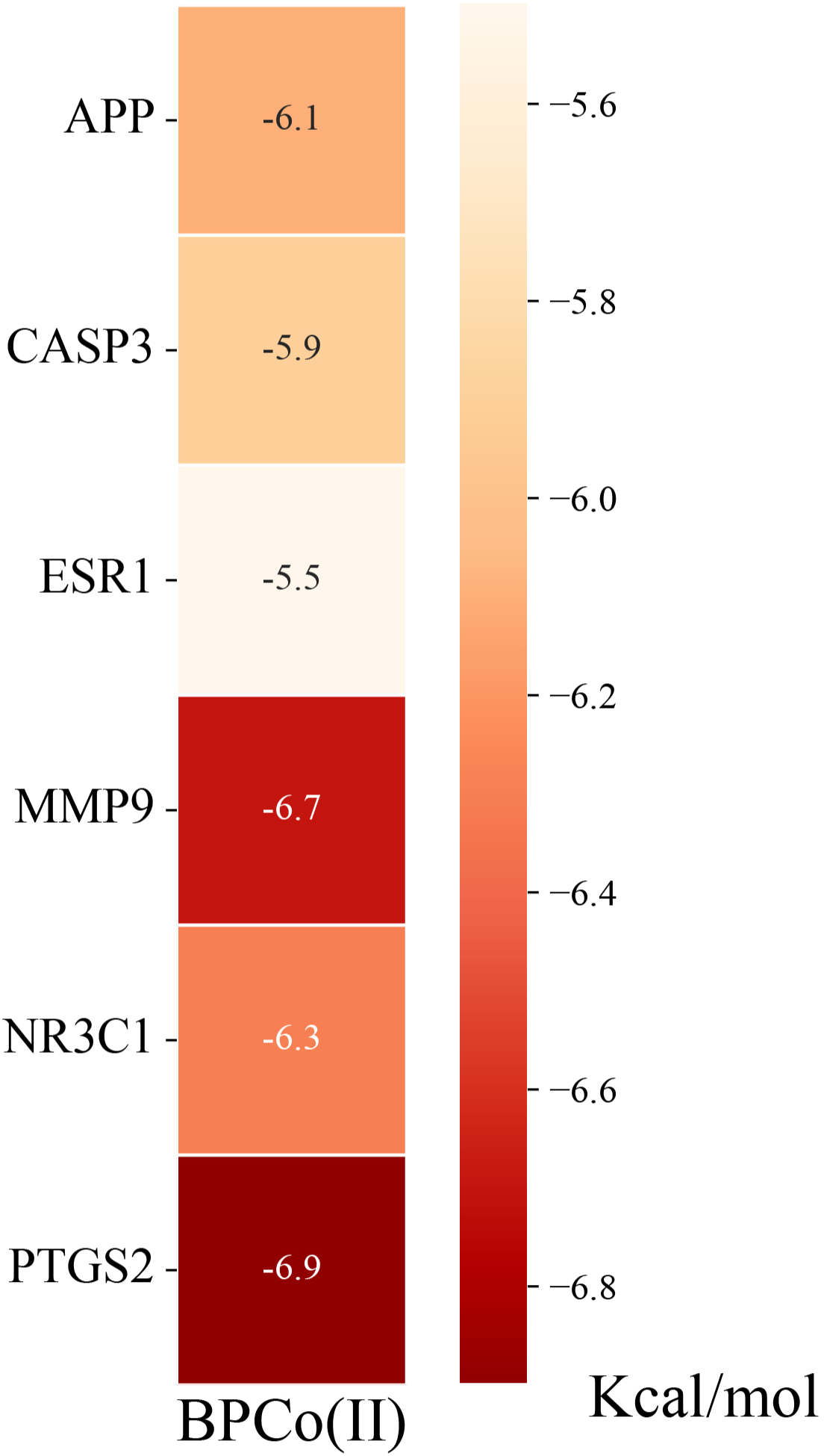
Heatmap of molecular docking binding energy of BPCo(II) to hub targets.

